# Opioid receptor distribution in the claustrum-dorsal endopiriform complex

**DOI:** 10.1101/2025.11.18.687877

**Authors:** Matthew Bolger, Jesse Jackson, Anna M.W. Taylor

## Abstract

The claustrum and dorsal endopiriform regions form a thin subcortical sheet reciprocally connected to the neocortex that is enriched in opioid receptors. The precise cellular distribution of opioid receptors within this region remains unclear. Using multiplexed fluorescence *in situ* hybridization (mFISH) and hierarchical cluster analysis, we mapped the expression of mu (*Oprm*), delta (*Oprd*), kappa (*Oprk*), and ORL1 (*Oprl*) opioid receptor genes at single-cell resolution in the mouse claustrum-dorsal endopiriform. Among 2,269 neurons analyzed, six transcriptional clusters were identified: three excitatory (CLA, CLA/OPRK, cortical excitatory) and three inhibitory (SST, PVALB, broad inhibitory). *Oprk* expression was uniquely restricted to excitatory claustrum neurons and particularly enriched in the CLA/OPRK cluster which was comprised largely of *Synpr*+ claustrum core projection cells. In contrast, *Oprd, Oprm*, and *Oprl* genes were more broadly distributed across both excitatory and inhibitory populations, with *Oprd* enriched in parvalbumin (*Pvalb)* inhibitory neurons and *Oprl* in somatostatin (*Sst)* inhibitory neurons. Spatial mapping confirmed that *Oprk* expressing cells were concentrated within the claustrum core, whereas other receptor subtypes extended across the entire claustrum and into adjacent cortical regions. A comparable distribution of opioid receptors was observed in the neighbouring dorsal endopiriform. Analysis of a publicly available single cell sequencing dataset of the macaque claustrum also revealed a similar receptor distribution, where *Oprk* expression was enriched within a subset of excitatory projection neurons, and *Oprm,Oprd*, and *Oprl* expressed widely in both inhibitory and excitatory cells. Together, these findings demonstrate an evolutionarily conserved, cell-type-specific organization of opioid receptor expression in the claustrum-dorsal endopiriform. These results indicate that kappa opioid receptor expression is a defining molecular feature of excitatory claustrum projection neurons and suggest distinct roles for other opioid receptors in modulating inhibitory and cortical circuits.

---

The claustrum and dorsal endopiriform regions comprise a thin sheet of subcortical grey matter that sits between the putamen and insula. This region spans the rostro-caudal axis of the brain and is reciprocally connected to much of the neocortex. Activation of this pathway coordinates cortical activity during online and offline states, and has been linked to behaviours associated with opioid signaling including pain ^1–3^, opioid reinforcement/relapse^4,5^ and stress^6^. This region is enriched in opioid receptor expression; however, the exact cellular distribution of mu (MOR), delta (DOR), kappa (KOR), and opioid receptor like-1 (ORL-1) opioid receptors is unknown.

The claustrum and endopiriform are made up of excitatory projection cells and several populations of inhibitory cells defined by peptide content, including parvalbumin, neuropeptide Y and somatostatin. Excitatory projection cells are topographically organized based on cortical projection targets^7,8^ and neurochemical profiles^9–11^. While the identification of the rodent claustrum-dorsal endopiriform complex has been historically quite challenging due to the thin structure and lack of obvious anatomical borders from adjacent cortical regions, recent gene sequencing data have identified a number of genes (*Synpr, Nr4a2, Lxn*) that have been validated markers of these regions ^9,12–15^.

Early studies examining opioid receptor binding and gene/protein expression have reported particularly high expression of ORL-1 (*Oprl)* and KOR (*Oprk)* in the claustrum-dorsal endopiriform region, with more modest expression of DOR (*Oprd*) and MOR (*Oprm*)^16–21^. More recent single cell sequencing experiments in mice have described *Oprk* and *Oprm* expression in excitatory claustrum cells, with relatively lower expression of *Oprd*^5,10^. Electrophysiological recordings of opioid agonist-evoked activity in the claustrum found MOR, DOR and KOR agonists reduced activity in claustrum projection cells; however, only KOR agonists reduced monosynaptic transmission^22^. This difference in activity suggests a wider, more nuanced distribution of opioid receptors beyond excitatory cells. We do not yet know the precise cellular and spatial distribution of all four opioid gene transcripts across all neuronal populations within the claustrum-dorsal endopiriform region.

Using multiplexed fluorescence *in situ* hybridization (mFISH) we sought to determine the distribution of the four opioid receptors at a single cell resolution in the claustrum-dorsal endopiriform. We then applied hierarchical cluster analysis to describe the expression of opioid receptors within excitatory and inhibitory clusters in the claustrum and dorsal endopiriform. Using a publicly available data set of claustrum gene expression in macaques, we determined the opioid expression pattern was similar between mice and non-human primates. Overall, these data indicate a distinctive and evolutionarily conserved pattern of expression of opioid receptors across the claustrum-dorsal endopiriform complex.

## Methods

All animal experiments and procedures were performed in compliance with the Canadian Council on Animal Care’s Guidelines and Policies with approval from the University of Alberta Health Sciences Animal Care and Use Committee. Four 23-week-old (2 male and 2 female) C57B16/J mice were euthanized using sodium pentobarbitol (Euthasol). Following a transcardiac perfusion of ice-cold saline, brains were taken and immediately immersed in cryo-embedding medium (OCT). After being cooled over dry ice, brains were moved to a -80°C freezer for storage.

Brains were sectioned at a thickness of 12⎧m using a cryostat tissue slicer and mounted on glass slides. mFish assay was performed as per RNAscope HiPlex12 v2 assay user manual (Advanced Cell Diagnostics) starting with the fresh frozen tissue pretreatment. Probes for mFish were purchased from Advanced Cell Diagnostics and include: *Nr4a2* (423351-T1), *Pvalb* (421931-T2), *GAD1* (400951-T3), *Lxn* (585801-T4), *Oprd* (427371-T5), *Oprm* (315841-T6), *Npy* (313321-T7), *Oprl* (514301-T8), *Sst* (404631 - T9), *Camk2a* (445231-T10), *Oprk* (316111-T11), *Synpr* (500961-T12). All 12 probes with unique tails (T1-T12) were hybridized to the tissue and following 3 successive amplification steps the first set of cleavable fluorophores were hybridized (T1-T4). Tissue was then counterstained with DAPI and coverslipped with ProLong Gold antifade mounting medium. Following image acquisition slides were decoverslipped by soaking in a saline-sodium citrate buffer and fluorophores (T1-T4) were cleaved. This process was repeated for 2 more rounds using fluorophores (T5-T8) and (T9-T12).

After each round of hybridization, Z-stack images (0.5 ⎧m step size) were captured using a Leica THUNDER-Deconvolution Widefield Microscope using a 63x oil immersion objective. During the first imaging session the claustrum-endopiriform complex region was identified through the expression of both *Nr4a2* and *Lxn*. For ease of analysis, the claustrum and dorsal endopiriform were captured separately. 4-9 contiguous images of the claustrum or dorsal endopiriform regions were captured and stitched together to form a single composite image. Maximum intensity projections from each round were loaded into Fiji^23^ and merged using the HyperStackReg plugin^24^. Merged images were then loaded into Qupath ^25^ to be pseudo colored and analyzed. Cell segmentation was performed on the DAPI channel of each merged image with the StarDist2D Qupath plugin using the pretrained model “dsb2018_heavy_augment.pb”^26^. The following parameters were used: probability threshold: 0.67, cell expansion: 5, cell constrain scale: 2, and pixel size: 0.5. This defined each cell’s nucleus and approximated the cytosol as a region of interest within each image. Probe expression was then detected within each region of interest utilizing the Qupath Subcellular detection function. Appropriate intensity thresholds were determined for each channel to optimize detection within a given image. The total area within each region of interest occupied by the signal of each probe was then quantified. Cell/probe detection results from each merged image were combined and exported to R (R Development Core Team, 2008) for computational analysis.

6,155 nuclei were registered across all mFish images of the claustrum-dorsal endopiriform complex. Probe expression was normalized using a probe’s percent area coverage (PAC) of its region of interest (the cell)^27^. Linear dimensionality reduction (PCA) was performed and principal components representing approximately 95% of variance were extracted and used for nonlinear dimensionality reduction (UMAP) with the umap package (Manhattan method, all other parameters default). From this, a large cluster of cells emerged which displayed little neuronal gene expression. Cells from this cluster were excluded and the remaining 2,269 cells were re-clustered. Hierarchical clustering (ward.D2 method) yielded strong agreement with UMAP dimensionality reduction, and UMAP clusters were colored based on hierarchical clusters.

Box plots were generated to visualize the distribution of gene expression in cells of a given cluster. Boxes represent the interquartile range with the center line being the median PAC for a given marker. Whiskers show distribution to at most 1.5x the interquartile range and outlier points are not shown. To more effectively contrast the level of gene expression between clusters, the z-score of each markers’ log(x+1) normalized PAC within a given cell was calculated. These standardized values were used to colour points on a gradient scale according to their level of gene expression in both UMAP and spatial plots. The Pearson coefficient was calculated for each pair of transcripts across all detected neurons to determine the degree of colocalization between opioid receptors and across inhibitory and excitatory clusters.

To assess the spatial distribution of different cell types, spatial plots were generated from the data of 5-6 images merged into a single plot. Images from both male and female animals were included in each merged plot. Each object (cell) generated through cell segmentation had their pixel coordinates along with marker expression levels and cluster affiliation recorded. As the images of the claustrum varied in size and orientation, location coordinates were transformed to be between 0 and 1 using the caret R package^28^. This allowed cells from different images to be plotted on the same axis while retaining their original spatial conformation. In a subset of plots, an ellipse generated from cells belonging to claustrum (CLA and CLA/OPRK) or dorsal endopiriform (dEpd/OPRK) related clusters were drawn at a confidence level of 0.8. An unpaired Wilcox test was used to investigate any potential sex differences in opioid receptor expression within a given cluster. Statistical significance was set at p<0.05.

## Results

### Classifying claustrum cells visualized by mFISH

mFISH was used to describe opioid receptor distribution in the claustrum-dorsal endopiriform complex at a single cell resolution. Several classical markers of excitatory and inhibitory cells were chosen to orient opioid receptor expression to specific subclasses of neurons. Calcium/calmodulin dependent protein kinase 2a (*Camk2a)* was used to identify broad excitatory cells, with nuclear receptor related protein 1 (Nurr1; gene *Nr4a2)*, latexin (*Lxn)*, and synaptoporin (*Synpr)* used as specific markers of claustrum-dorsal endorpiriform excitatory cells. Glutamate decarboxylase 1 (*Gad1)* was used to identify inhibitory cells, and parvalbumin (*Pvalb)*, neuropeptide Y (*Npy)* and somatostatin (*Sst)* used to demarcate specific inhibitory populations. The expression of these 12 genes was spatially registered across sections of the claustrum and dorsal endopiriform (Fig 1A). The percent area covered (PAC) by each fluorescently tagged probe in each cell was calculated to compare gene expression levels across cells.

**Figure 1:**
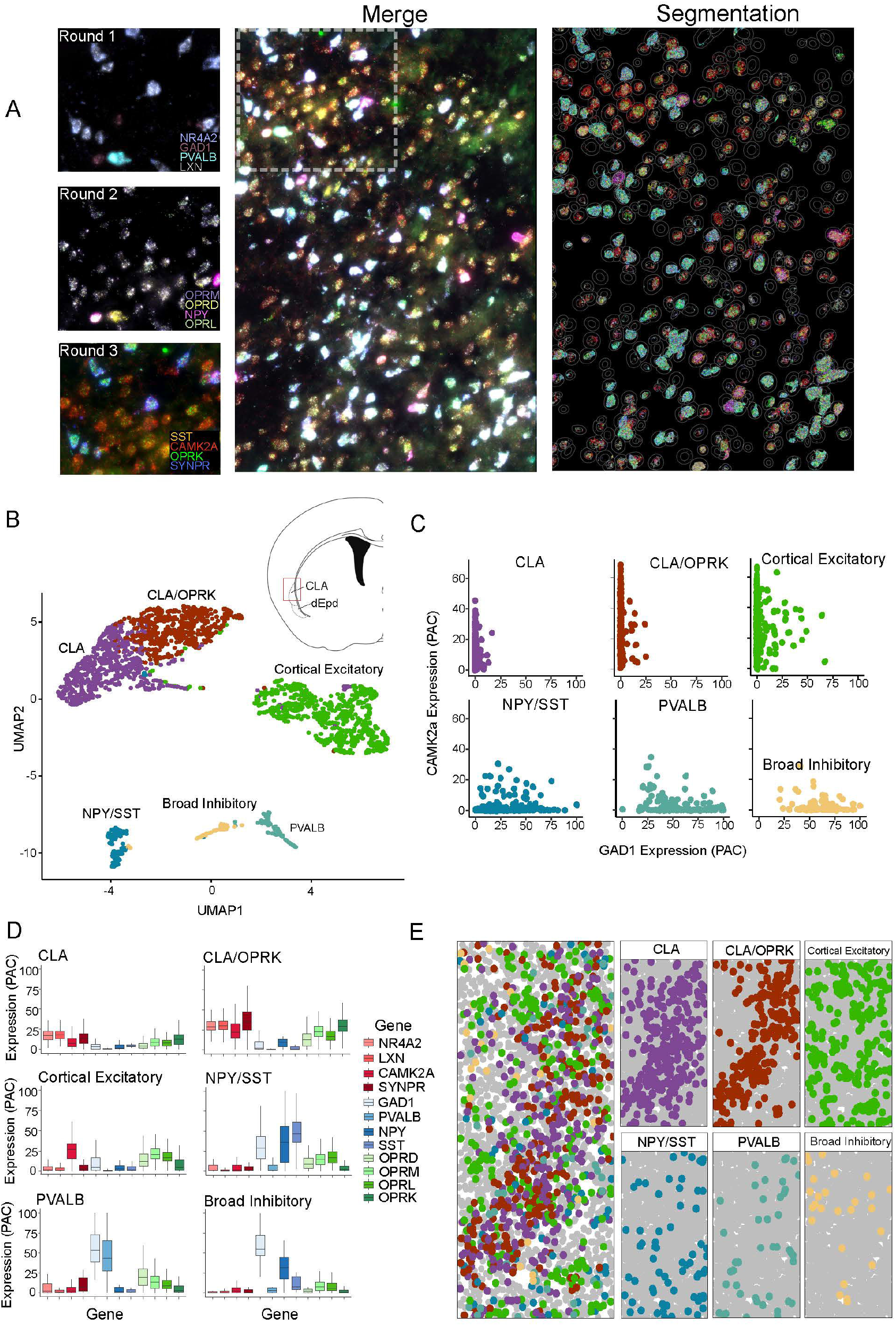
Identification of claustrum neuronal populations utilizing multiplexed fluorescent *in situ* hybridization. (A) Representative images of the claustrum. Labelling for all 12 probes occurred in 3 iterative steps on the same brain slice. After each step, images were acquired (left) and spatially registered images were merged into one 12-channel imaged (middle). A cell segmentation tool was used to define regions of interest that was used for analysis (right). Scale bars =50um (B) UMAP based non-linear dimensionality reduction of processed claustrum cell expression data, labelled and coloured as per hierarchical clustering assignment. (C) *Camk2a* expression plotted against *Gad1* expression for each identified cluster. PAC = percent area covered and indicates the fraction of cell occupied by fluorescently tagged gene probe (D) Expression levels of all genes for each identified claustrum cell cluster. (E) Cells identified in B plotted using scaled spatial coordinates. Scale bar = 50 um.

Through hierarchical clustering analysis, 6 patterns of marker co-expression were identified in the claustrum (Fig 1B). Three of the identified clusters were enriched with the excitatory gene *Camk2a* (CLA, CLA/OPRK, cortical excitatory), while three were enriched with the inhibitory gene *Gad1* (NPY/SST, PVALB, Broad Inhibitory). To further confirm this broad classification, the level of *Gad1* expression within each cluster was compared against the expression of *Camk2a* (Fig 1C). Clusters with high expression (over 50%) of *Camk2a* (CLA, CLA/OPRK, and Cortical Excitatory) had less than 1.5% coverage by *Gad1* (OPRK/CLA=0.38%, Cortical Excitatory =1.36%, CLA=0.21%). Conversely, clusters with high expression (over 75%) of *Gad1* had less than 4% coverage by *Camk2a* (NPY/SST=3.05%, Pvalb=3.74%, Broad Inhibitory=3.21%). This indicates that *Camk2a* and *Gad1* expression reliably identifies segregated populations of putative excitatory and inhibitory cells. Of the cells identified in the claustrum, approximately 85% belonged to excitatory clusters and 15% belonged to inhibitory clusters.

Two of the excitatory clusters prominently expressed all three claustrum markers (*Synpr, Nr4a2, Lxn*) indicating two distinct subpopulations of claustrum excitatory cells. However, one of these subpopulations exhibited higher marker expression across the board, with the two most differentially expressed markers being *Synpr* and *Oprk*. This population was classified as a CLA/OPRK cell cluster while the former was classified as the CLA cell cluster. These two excitatory claustrum clusters also exhibited notable spatial distributions, with the CLA/OPRK cells clustering towards the core of the claustrum and the CLA cells more widely distributed across the core and shell of the claustrum (Fig 1E). The third excitatory cluster did not significantly express any claustrum markers but expressed spatial and transcriptomic properties of cortical excitatory cells. As such it was classified as “cortical excitatory” and likely represents generalized excitatory cells expressed across the neocortex.

Of the cells classified as inhibitory (*Gad1*+), the largest cluster was enriched in *Sst* and *Npy*. The other major inhibitory cluster prominently expressed *Pvalb*. The final, smaller, population prominently expressed only *Gad1*, with *Npy* being expressed to a lesser extent. There were no identifiable spatial distribution patterns within the inhibitory cells likely due to a lack of cells compared to the excitatory clusters.

### Opioid receptor expression varies by claustrum cell type

We next assessed the expression of all four opioid receptors within the identified clusters (Fig 2A, B). KOR (*Oprk)* exhibited a distribution within the observed cell populations distinct from the other opioid receptors. *Oprk*, was most prominently expressed within the excitatory CLA/OPRK cluster. *Oprk* expression was detectable in other excitatory clusters (CLA and Broad Excitatory), but at significantly lower levels than the CLA/OPRK cluster. Notably, *Oprk* expression was not detectable above background in any of the inhibitory clusters. In contrast, expression of the other opioid receptor genes exhibited much wider distribution across both inhibitory and excitatory clusters. *Oprd* expression was highest in the PVALB inhibitory cluster; however, it was also present consistently within the Cortical Excitatory, CLA/OPRK and NPY/SST cell populations. *Oprl* expression was highest in the NPY/SST cluster; however, it was also present consistently within the cortical excitatory and CLA/OPRK clusters. *Oprm* expression was also found in both excitatory and inhibitory clusters but expressed at higher levels within most excitatory populations. Apart from *Oprk* and the CLA/OPRK cluster, no other opioid receptor gene was sufficient to define a specific cluster or cell type.

**Figure 2:**
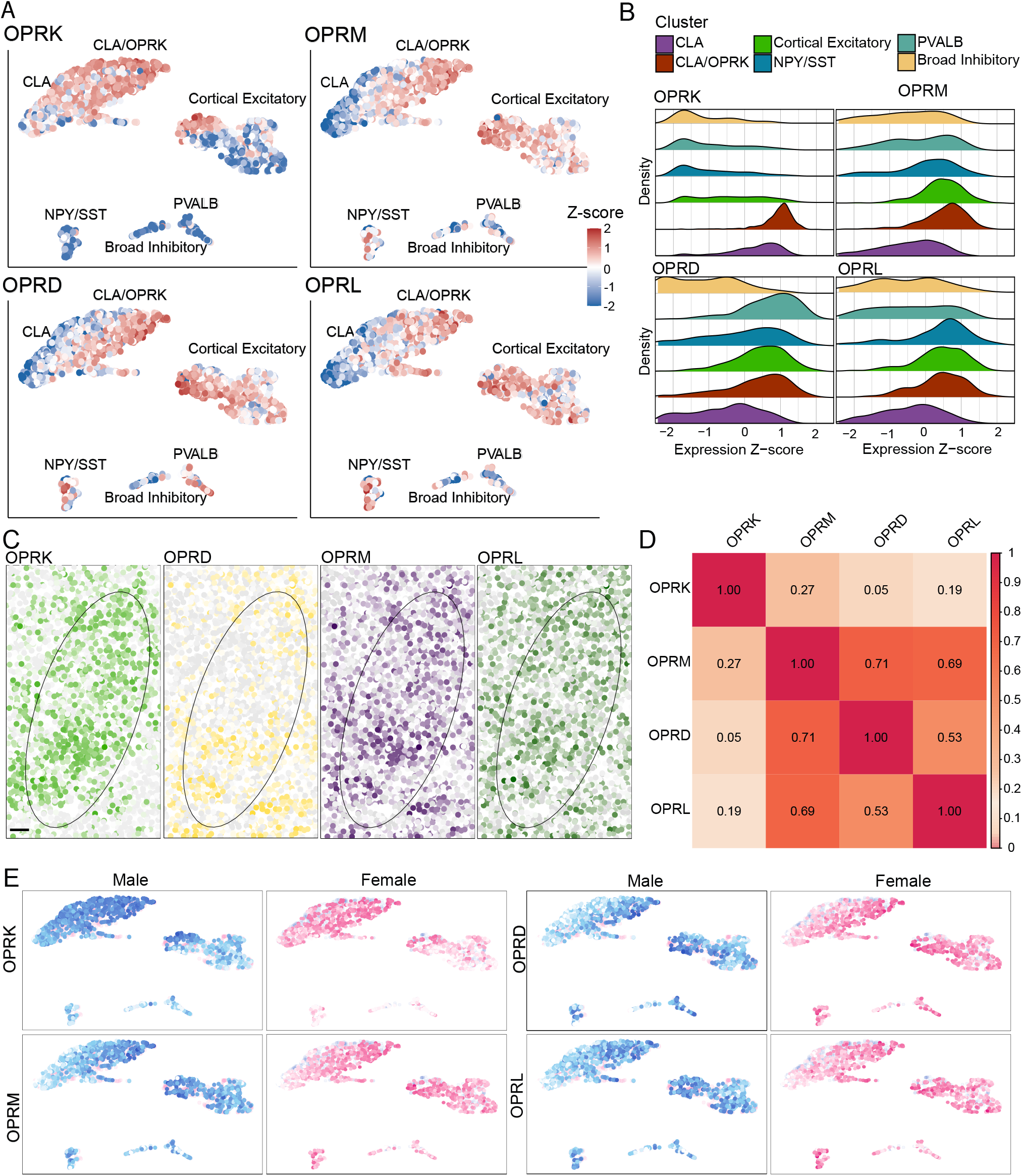
Opioid receptor expression within the claustrum. (A) Labelled UMAP projection coloured as per each cell’s expression of each opioid receptor individually. (B) Stacked density plots of opioid receptor expression within each identified cluster. (C) Spatial plot as in 1(E) but coloured as per each cell’s z-score of opioid receptor expression. Claustrum location approximated based on the position of putative claustrum cells (CLA and CLA/OPRK clusters) and indicated with an ellipse indicating the region covered by 80% of the CLA-specific excitatory cells. Scale: Range from -2 to +2, Coloured from grey to assigned colour. (D) Pearson correlation table for the opioid receptor expression across all claustrum cells. (E) UMAP of opioid receptor expression from male and female brains. Cells coloured as per sex of animal, blue (male) or pink (female). Scale: Range from -2 to +2, Coloured from white to assigned colour, Cells of opposing sex present but with reduced opacity.

Of all the opioid receptor genes, *Oprk* was most restricted to the region identified as the claustrum (defined by *Nr4a2* expression, Fig 2C). Cells expressing *Oprd, Oprm, or Oprl* showed a wider spatial distribution that extended beyond the claustrum into the neighbouring cortex. *Oprk* expression was the least correlated with other opioid receptor genes (Fig 2D, Fig S1), indicating that cells expressing *Oprk* are less likely to also express other opioid receptor genes. In contrast *Oprm, Oprl, and Oprd* expression were more consistently correlated with each other, with *Oprm* expression showing the highest degree of correlation with the other opioid receptors. This indicates that cells that express *Oprm* are more likely to express at least one other opioid receptor gene (besides *Oprk)*.

Finally, opioid receptor distribution in the claustrum was compared between male and female tissue (Fig 2E, Fig S2). No significant differences in opioid receptor expression were noted in the majority of cell populations. The only exception was *Oprd* expression within the CLA/OPRK population, which was higher in females than males.

### Dorsal endopiriform cell populations and opioid receptor distribution is similar to those of the claustrum

The dorsal endopiriform is adjacent to the claustrum and shares common anatomical and neurochemical features. This region has been defined as the extended claustrum and can be identified based on expression of Nurr1 (*Nr4a2*)^12^. Here, we compared the cellular distribution of opioid receptors within the dorsal endopiriform and compared this to the distribution in the claustrum (Fig 3). Using the same data analysis pipeline as before, we found 585 cells with significant neuronal marker expression. These cells formed 4 clusters which were similar to those populations defined in the claustrum (Fig 3A, B). The first identified cluster was an excitatory cluster which showed high expression of *Nr4a2, Lxn, Synpr*, and *Oprk*. This population was defined as the dEPd/OPRK cluster and was analogous to the CLA/OPRK population defined in the claustrum. While *Oprk* expression within this dorsal endopiriform population was comparatively lower than the claustrum, these cells still represented the primary *Oprk*+ cell population. Like their claustrum analogues, this cluster was spatially distributed towards the core of the dorsal endopiriform (Fig 3C).

**Figure 3:**
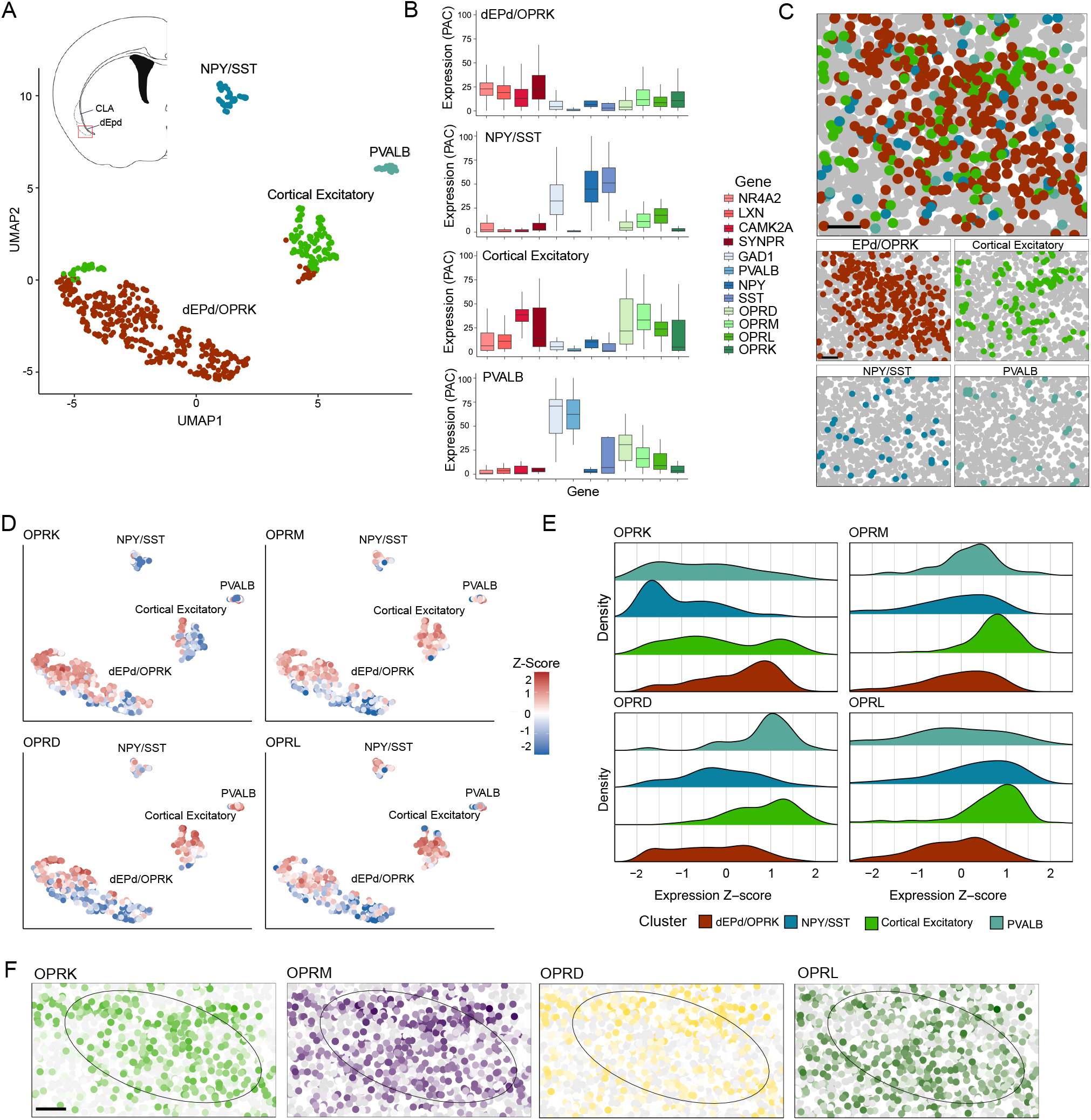
Opioid receptor expression in the dorsal endopiriform. (A) UMAP based non-linear dimensionality reduction of processed endopiriform cell expression data, labelled and coloured as per assigned cluster. (B) Expression levels of all genes for each identified endopiriform cell cluster. (C) Cells identified from A plotted using scaled spatial coordinates. (D) Labelled UMAP projection coloured as per each cell’s expression of each opioid receptor individually. (E) Stacked density plots of opioid receptor expression within each identified cluster. (F) Spatial plot as in (C) but coloured as per each cell’s z-score of opioid receptor expression. Endopiriform location approximated based on position of excitatory dorsal endopiriform cells (dEPd/OPRK cluster) and indicated with an ellipse indicated 80% of the region covered by dorsal endopiriform excitatory cells. Scale: Range from -2 to +2, Coloured from grey to assigned colour.

The next identified cluster was also deemed excitatory due to the prominent expression of *Camk2a* (Cortical Excitatory). The cells within this cluster expressed relatively less *Nr4a2* and *Lxn* and therefore lack defined markers for the claustrum-dorsal endopiriform region. This suggests this population represents a broad cortical excitatory population analogous to the Cortical Excitatory population defined in the claustrum. This identity is further supported by the pattern of opioid receptor expression, in which it is enriched in *Oprm, Oprd*, and *Oprl* genes with relatively lower levels of *Oprk* expression (Fig 3D, E).

The remaining two clusters were classified as inhibitory based on their high levels of *Gad1*. The NPY/SST cluster was notable for its high expression of *Npy* and *Sst* (Fig 3 A,B). Much like in the claustrum, this cluster was enriched with *Oprl*, while *Oprk* was largely unexpressed (Fig 3D, E). The PVALB cluster was defined by the high level of *Pvalb* expression, and consistent with the claustrum, was enriched with *Oprd*. No distinguishable spatial distribution was observed (Fig 3C). There were no notable sex differences in opioid receptor expression within any dorsal endoprirform cell population (Fig S3).

Overall, the opioid receptor distribution in the dorsal endopiriform was consistent with the patters observed in the claustrum. *Oprk* expression was largely limited to excitatory cells that express claustrum-dorsal endopiriform markers such as *Nr4a2, Lxn* and *Synpr*. Meanwhile the other opioid receptor subtypes showed no such specificity for excitatory cells. Like in the claustrum, high expressing *Oprk* cells tended to be concentrated together in the core region of the dorsal endopiriform. No other opioid receptor subtype showed any identifiable spatial trends.

### Macaque Claustrum

The claustrum is a highly conserved structure with analogous regions in the human and non-human primate^11,29^. Using a publicly available single cell sequencing dataset of the macaque claustrum^11^, we described the distribution of all four opioid receptors within defined clusters of both neuronal and non-neuronal cells. *Oprk* expression was enriched in a subpopulation of excitatory cells (Fig 4). This cluster was defined by expression of *Gnb4*, which represented the largest excitatory population in the macaque claustrum. *Oprk* expressing cells accounted for approximately a third of the total population of this *Gnb4* cluster and was only lowly expressed within inhibitory clusters.

**Figure 4:**
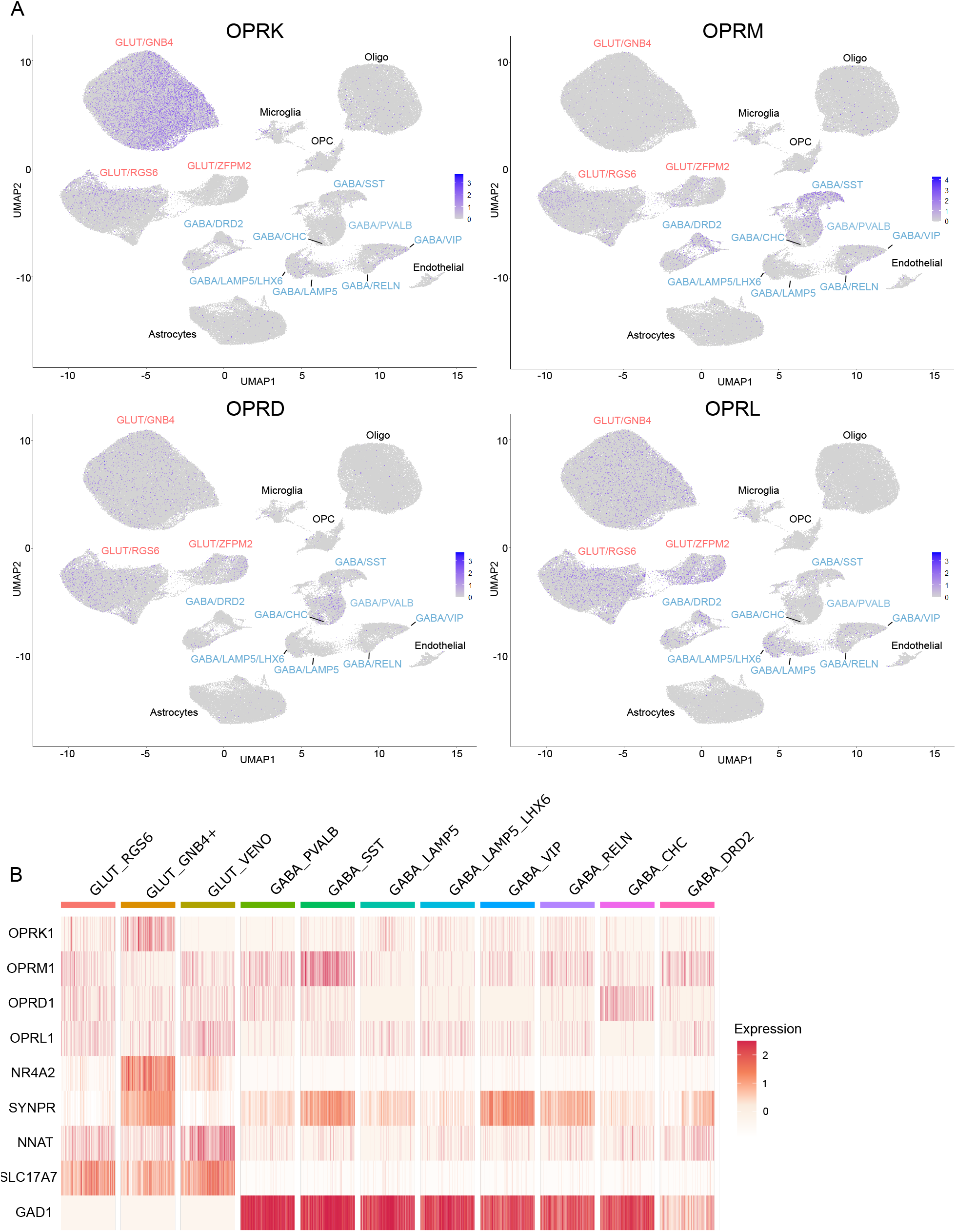
Opioid receptor expression in the macaque claustrum. **(A)** UMAP based representation opioid receptor expression within all identified macaque claustrum cell populations. Cells coloured based on level of expression represented through pearson residuals. Excitatory clusters labelled in red, inhibitory clusters labelled in blue, non-neuronal cells labelled in black (B) Gene expression heatmap of both opioid receptors and select population markers within identified neuronal populations down sampled to 1000 cells.

Like the mouse, the expression of the other opioid receptors was less population specific. Interestingly, *Oprm* expression was prominently seen within inhibitory populations, especially the *Sst* subpopulations. Within excitatory cells, *Oprm* was most prominent within the population of deep layer 6 insular cells (GLUT/ZFPM2). *Oprd* and *Oprl* expression within excitatory cells was similarly biased towards this population, although they were more widely expression across all excitatory populations than *Oprm*. Within inhibitory populations, *Oprd* was preferentially expressed in parvalbumin (PVALB/UNC5B) and chandelier (CHC) inhibitory neurons. Meanwhile *Oprl* was expressed across most of the cell populations at consistent levels, with parvalbumin and CHC inhibitory neurons the only where *Oprl* was largely absent. Overall, these results suggest that opioid receptor distribution is largely conserved across species.

## Discussion

The claustrum-dorsal endopiriform complex is a region enriched in opioid receptors. However, the exact cellular and spatial distribution of all four opioid receptors was unknown. Here, we used multiplexed fluorescent *in situ* hybridization and spatial transcriptomics to describe the pattern of expression of MOR, DOR, KOR, and ORL1 receptors in the claustrum-dorsal endopiriform complex.

In line with previous gene and protein expression studies, all four opioid receptors were detected with the claustrum and dorsal endopiriform. *Oprk* expression was distinctive in that it was the only receptor that was restricted to excitatory neurons. *Oprk* expression was highest in clusters containing defined markers for claustrum cells, and lower in the neighbouring cortical clusters. This is in line with other studies that indicates KOR expression is highest in the claustrum relative to neighbouring cortical regions^5,11^. Previous studies have identified two spatially distinct projection cell populations within the claustrum termed “core” and “shell”, with core cells projecting to frontal/midline cortical structures and shell cells projecting preferentially to posterior structures^7–10^. Core claustrum projection cells can be identified based on their high expression of *Nr4a2, Lxn*, and *Synpr* ^9^. Here, we found *Oprk* was most highly expressed within the claustrum excitatory cluster enriched with *Synpr* (CLA/OPRK cluster). This cluster had relatively higher levels of *Nr4a2* and *Lxn* than the other claustrum excitatory cluster (CLA), suggesting *Oprk* expression is most highly expressed on claustrum core projection cells. This was confirmed with spatial analysis that found CLA/OPRK cells clustered towards to the core of the claustrum. The expression of *Oprk* in core claustrum projection cells is supported by several other lines of evidence. For example, previous sequencing experiments have reported that *Oprk* is enriched in a single excitatory population^5,10^. Nurr-1 (*Nr4a2)* is a key regulator guiding the development of the claustrum core cell molecular phenotype^10^. When Nurr-1 was knocked down in the claustrum, *Oprk* expression was significantly downregulated^12^. While a nuanced classification of dorsal endopiriform projection cells based on neurochemical or anatomical projection targets have not yet been described, our spatial transcriptomics assessment reveals similar distribution of *Oprk* to the claustrum, where it was enriched in the excitatory endopiriform cluster (dEpd/OPRK). Overall, these results indicate KOR receptors are expressed on a subpopulation of core claustrum-dorsal endopiriform excitatory cells. While the functional ramifications of this distribution remain to be studied, the preferential expression on core cells suggest KOR ligands will more strongly influence claustrum output modules targeting frontal/midline cortical regions.

*Oprd, Oprl*, and *Oprm* exhibited a much wider distribution across the excitatory and inhibitory clusters within the claustrum and dorsal endopiriform. This has been shown to have functional implication for how opioid agonists impact claustrum function. For example, while KOR agonists abolished optogenetically evoked excitatory activity in the claustrum, MOR and DOR agonists only reduced slower, long latency recurrent excitatory responses^22^. While *Oprd* and *Oprl* expression was similarly expressed within all excitatory populations, these genes showed dissociable patterns of distribution within the inhibitory subpopulations. While *Oprd* expression was highest in the *Pvalb* inhibitory cluster (PVALB), *Oprl* expression was highest within the *Sst* inhibitory cluster (NPY/SST). Similar to the spatial distribution of claustrum excitatory neurons described above, inhibitory populations are also arranged in a mosaic core/shell topography with *Sst* inhibitory cells located within the claustrum shell and *Pvalb* inhibitory cells within the claustrum core^8^. This indicates that *Oprd*, located preferentially on *Pvalb+* cells, will have a stronger disinhibitory impact on core claustrum cells that project to frontal midline regions over neurons in the claustrum shell projecting to posterior cortical regions. In contrast, *Oprl* expression was highest within the *Sst* population that indicates it will preferentially impact shell projection cells. The functional impact of this differential expression of *Oprd* and *Oprl* on inhibitory populations remains to be determined.

Finally, we examined whether opioid receptor expression in the claustrum-dorsal endorpiriform region was conserved across species. Using a publicly available dataset of single cell sequencing of the macaque claustrum, we found a similar distribution of *Oprk* expression restricted to a subpopulation of claustrum projection cells (*Gnb4+*)^11^. Interestingly, only approximately 30% of this excitatory cluster expressed *Oprk*, compared to the mouse claustrum where approximately 70% express *Oprk*. The relative reduction in *Oprk* expression coincided with an overall expansion of the non-*Oprk+ Gnb4* cluster in macaques and marmosets, which suggests an expansion of the functional roles of *Gnb4*+ cell population in primates. Like in the mouse, the other opioid receptors exhibited a wider distribution across most inhibitory and excitatory populations, with no obvious cellular or spatial distribution.

Overall, these data show a conserved, cell-type-specific organization of opioid receptors across the claustrum-dorsal endopiriform complex. All four opioid receptors were highly expressed within the region, and most cell populations expressed more than one opioid receptor. Notably, only the KOR gene, *Oprk*, emerged as a defining molecular feature of excitatory claustrum core projection neurons, indicating KOR signalling will selectively influence a subset of claustrum output modules targeting frontal cortical regions.

## Figures Legends

**Figure S1: Correlation matrix for all genes.** Pearson correlation table for expression of all 12 transcripts across clustered claustrum cells.

**Figure S2: Opioid receptor expression in the male and female claustrum.** Box plots comparing percent area coverage (PAC) of each opioid receptor gene within each cell cluster. Unpaired Wilcox test was used to compare expression levels between males and females. ns = not significant, *=p<0.05.

**Figure S3: Opioid receptor expression in the male and female dorsal endopiriform. A)** Expression levels of opioid receptors present in each endopiriform cell cluster within both male and female images. B) Box plots comparing percent area coverage (PAC) of each opioid receptor gene within each cell cluster. Unpaired Wilcox test was used to compare expression levels between males and females. ns = not significant.

## Acknowledgements

The authors would like to thank Mark Cembrowski for helpful discussions and technical guidance. AMWT is supported by the Alberta Cancer Foundation Chair in Palliative Care and a Canada Research Chair Tier 2 in Pain and Addiction. This work was supported by Natural Sciences and Engineering Research Council of Canada (RGPIN 2018-04010 to AMWT) and the Canadian Institutes for Health Research (PJT 192040 to AMWT; PJT 195864 to JJ). Microscopy work was supported through the Cell Imaging Core at the University of Alberta.

