## Supplementary figures and images for "Opioid receptor distribution in the claustrum-dorsal endopiriform complex"

### Supplemental Figure 1

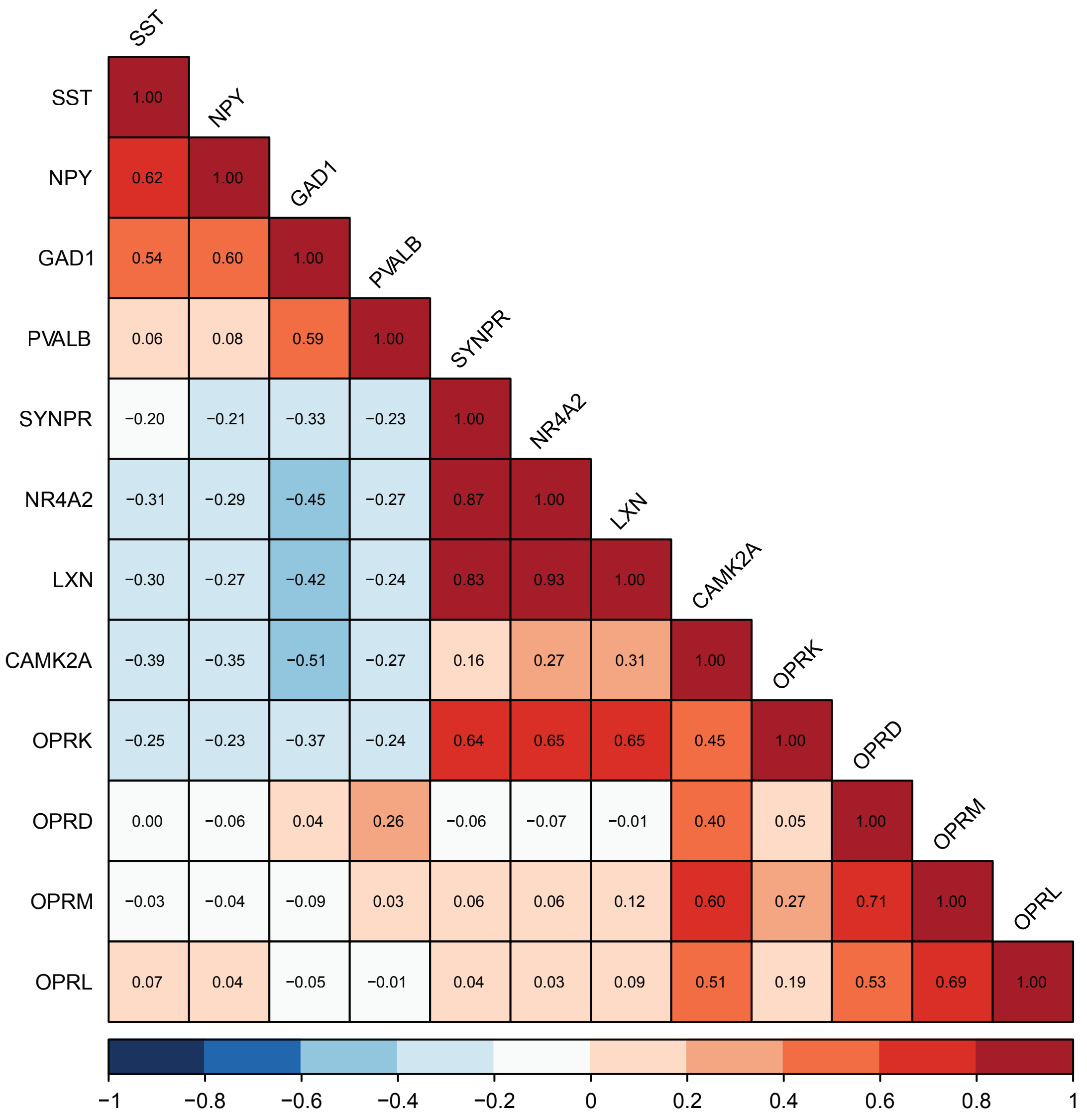

### Supplemental Figure 2

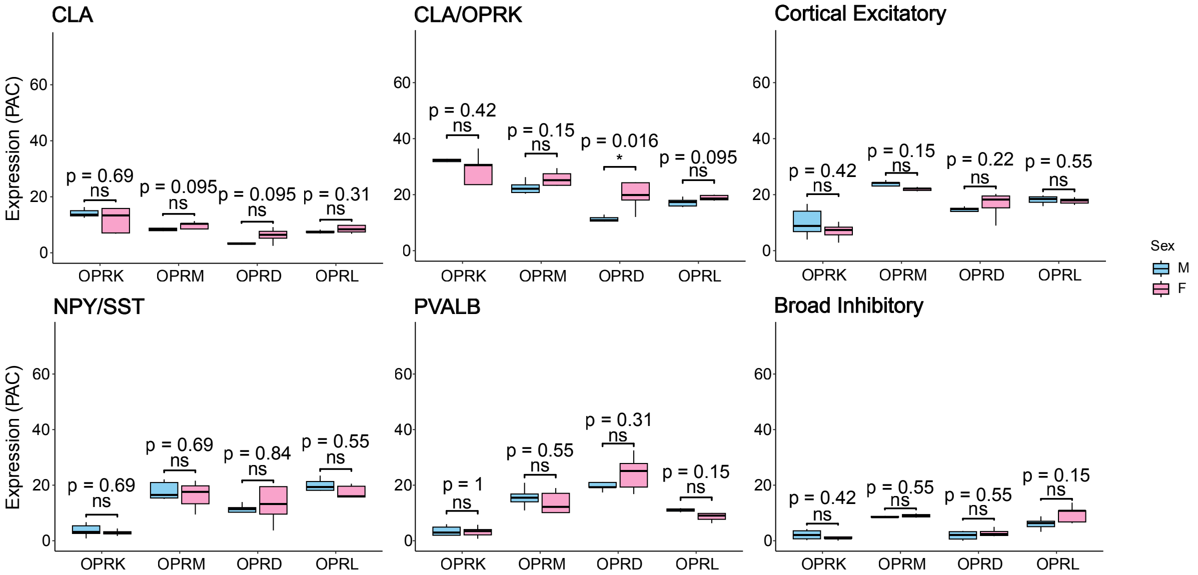

### Supplemental Figure 3

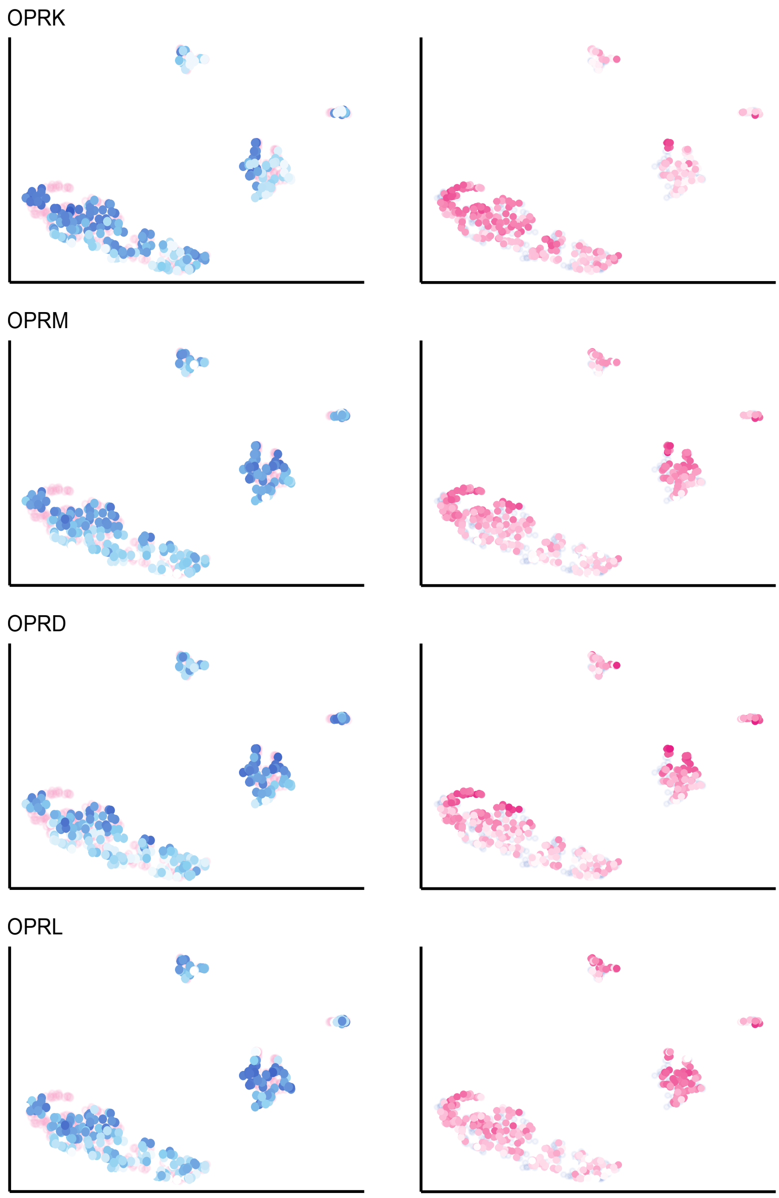

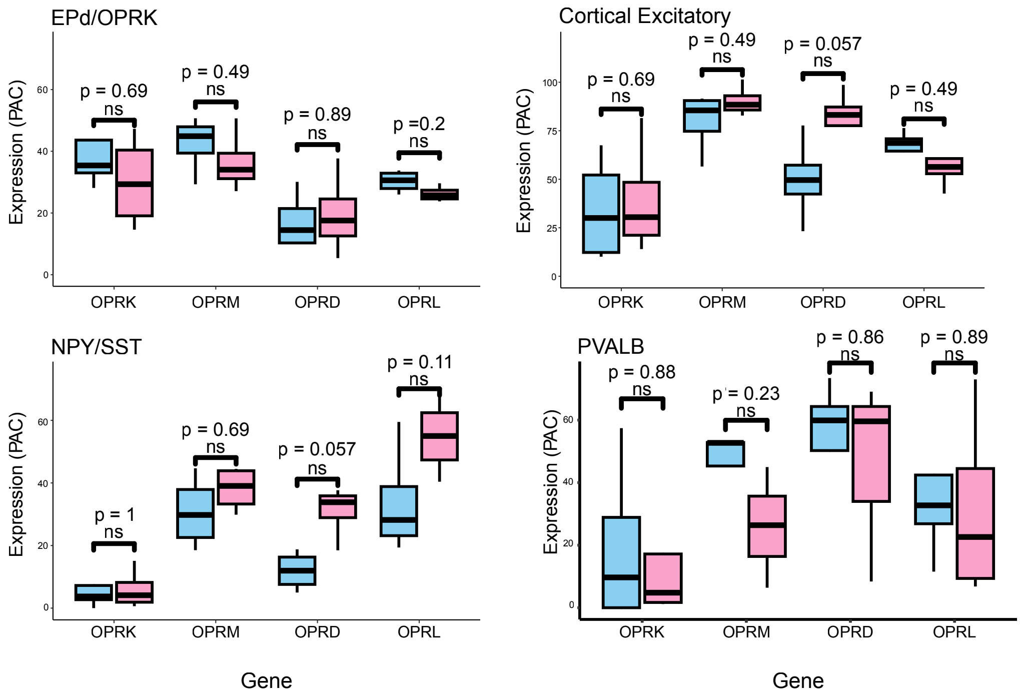
